# T2T-YAO: a Telomere-to-telomere Assembled Diploid Reference Genome for Han Chinese

**DOI:** 10.1101/2023.07.17.549286

**Authors:** Yukun He, Yanan Chu, Shuming Guo, Jiang Hu, Ran Li, Yali Zheng, Xinqian Ma, Zhenglin Du, Lili Zhao, Wenyi Yu, Jianbo Xue, Wenjie Bian, Feifei Yang, Xi Chen, Pingan Zhang, Rihan Wu, Yifan Ma, Changjun Shao, Jing Chen, Jian Wang, Jiwei Li, Jing Wu, Xiaoyi Hu, Qiuyue Long, Mingzheng Jiang, Hongli Ye, Shixu Song, Guangyao Li, Yue We, Yu Xu, Yanliang Ma, Yanwen Chen, Keqiang Wang, Jing Bao, Wen Xi, Fang Wang, Wentao Ni, Moqin Zhang, Yan Yu, Shengnan Li, Yu Kang, Zhancheng Gao

**Affiliations:** Department of Respiratory and critical care medicine, Peking University People’s hospital, Beijing, 100044, China; Beijing Institute of Genomics, Chinese Academy of Sciences/China National Center for Bioinformation, Beijing, 100101, China; Linfen Clinical Medical Research Center, Linfen, 041000, China; Grandomics Biosciences, Beijing, 102200, China; Department of Respiratory, Critical Care and Sleep Medicin, Xiang’an Hospital of Xiamen University. School of Medicine, Xiamen University, Xiamen, China; Institute of PSI Genomics, Wenzhou, 325024, China; Beijing Jishuitan Hospital, Capital Medical University, Beijing, China; Peking University Health Science Center, Beijing, 100191, China; Beijing Key Laboratory of Genome and Precision Medicine Technologies, Beijing, 100101, China; Institute of Chest and Lung Diseases, Shanxi Medical University, Taiyuan 030001, China

**Author notes:** These authors contribute equally (Yukun He, Yanan Chu, Shuming Guo, Jiang Hu, Ran Li, Yali Zheng). Corresponding authors: Zhancheng Gao, Department of Respiratory and critical care medicine, Peking University People’s Hospital, Beijing, 100044, China;., Yu Kang, CAS Key Laboratory of Genome Sciences and Information, Beijing Institute of Genomics, Chinese Academy of Sciences, 100101, Beijing, PR China.

**Keywords:** Reference genome, telomere-to-telomere assembly, Han Chinese, haplotype-resolved assembly, genome polishing, quality value, structure variation, small variation, genome annotation, exclusive gene

## Abstract

Since its initial release in 2001, the human reference genome has been continuously improved in both continuity and accuracy, and the recently-released telomere-to-telomere version—T2T-CHM13—reaches its top quality after 20 years of effort. However, T2T-CHM13 does not represent an authentic diploid human genome, but rather one derived from a simplified, nearly homozygous genome of a hydatidiform mole cell line. To address this limitation and provide an alternative pertinent to the Chinese population, the largest ethnic group in the world, we have assembled a complete diploid human genome of a male Han Chinese, T2T-YAO, which includes telomere-to-telomere assemblies for all the 22+X+M and 22+Y chromosomes in his two haploids inherited separately from his parents. Both haplotypes contain no artificial sequences or model nucleotides and possess a high quality comparable to CHM13, with fewer than one error per ∼14 Mb. Derived from the individual who lives in the aboriginal region of Han Chinese, T2T-YAO shows clear ancestry and potential genetic continuity from the ancient ancestors of the Han population. Each haplotype of T2T-YAO possesses ∼340 Mb exclusive sequences and ∼3100 unique genes as compared to CHM13, and their genome sequences show greater genetic distance to CHM13 than to each other in terms of nucleotide polymorphism and structural variations. The construction of T2T-YAO would serve as a high-quality diploid reference that enables precise delineation of genomic variations in a haplotype-sensitive manner, which could advance our understandings in human evolution, hereditability of diseases and phenotypes, especially within the context of the unique variations of the Chinese population.

## MAIN TEXT

A complete and accurate reference genome has been a long-standing goal in the biomedical research community since the initiation of the Human Genome Project three decades ago. However, the limitations of sequencing technology have made it challenging to achieve this level of completeness and accuracy(1–3). Recently, a groundbreaking publication from the Telomere-to-Telomere (T2T) Consortium described the first-ever complete haploid human genome of exceptional quality known as CHM13. This success has fulfilled 8% of the previously unknown highly repetitive region in human genome (4), which is attributed to significant improvements in sequencing technology and supporting bioinformatics tools. The resulting T2T-CHM13, an even more comprehensive reference for genetic research and various-omics studies, allows for more accurate localization of variation signals, especially in regions of repeats and duplications(5–10).

Despite the significant advancements in sequencing technology, assembling a high-quality diploid telomere-to-telomere (T2T) reference genome for an individual has yet to be achieved, even for the extensively sequenced and assembled diploid genome HG002(11). CHM13, while a remarkable scientific achievement, is a haploid genome of the complete hydatidiform mole (CHM) lacking the Y chromosome(12). Even by adding the complementary Y chromosome of HG002, the v2.0 of CHM13 is not enough for representing all individuals worldwide, as emphasized by the initial release of the Human Pangenome Reference Consortium (HPRC) which includes draft genomes from 47 individuals worldwide(13). The pangenome study reveals millions of single-nucleotide polymorphisms (SNPs), small indels, and tens of thousands of structural variations (SVs) per haplotype, with at least 100 Mb of sequences (∼3%) not represented in CHM13 which is of northern European origin except its Eastern European Ashkenazi Jewish ancestry chromosome Y (13). Similar findings are also reported in a recent pangenome study of Chinese populations (14), highlighting the importance of creating distinct reference genomes for each major population. Population-stratified references would allow for a more comprehensive understanding of genome variation across different populations for in-depth biological research and medical application.

The Han Chinese is the largest population in the world with billions of descendants worldwide, and yet remains underrepresented in current human reference genomes, including GRCh38 and the Human Pangenome Reference, especially lack of samples from their aboriginal regions. Despite extensive efforts being made to create reference genome for the Han population, including NH1.0 (15), HJ(16), Han1(17), and CN1(18), technical limitations make this task difficult for an accurate and complete one. Here, we have constructed a diploid T2T reference genome with a Y chromosome for the Han population (**Figure 1A**). Our sample comes from a healthy young man living in an ancient village in Shanxi province that has been inhabited by Han population for tens of generations as recorded in their family genealogy. We have named this reference genome “T2T-YAO” after the sampling point which is located near the ruin capital of the Emperor YAO from thousands of years ago. This area is also significant as it marks the starting point of the Hongtong migration during the Ming Dynasty. This migration, which lasted for nearly half a century (AD 1370 – 1417), saw large amount of emigrants throughout China and into Southeast Asia. Thus, the T2T-YAO genome is expected to provide a comprehensive representation of the Han population, and the reference-quality it achieves, *i.e.*, less than one error per ∼14 Mb sequences, enables its applications in future medical research and clinical practice for this vast population group.

**Figure 1.**
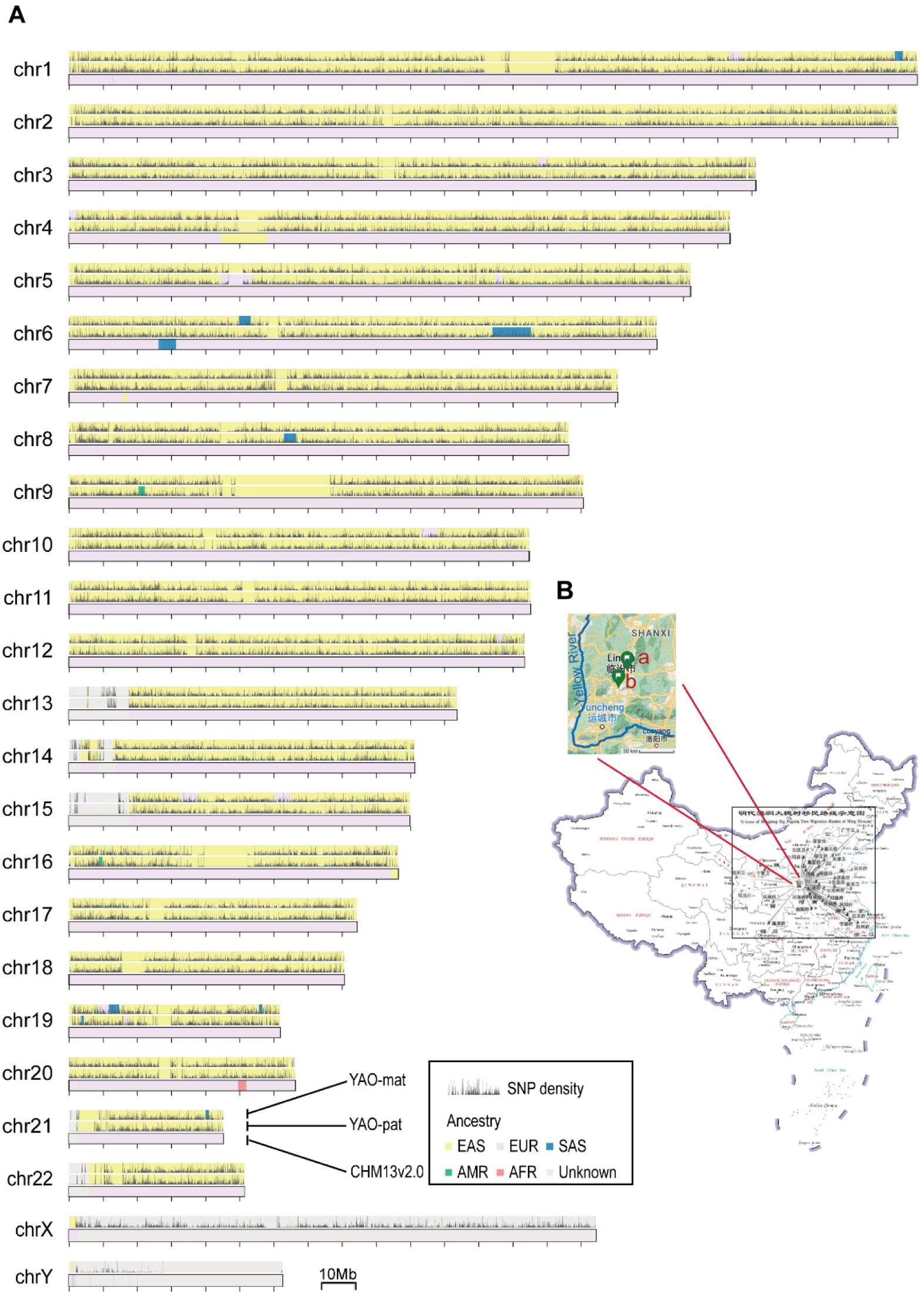
Overview of T2T-YAO and its ancestry. **A**. Comparison of the Ancestry markers of the T2T-YAO and CHM13(v2.0). Each set of chromosome includes maternal haplotype of Yao (top), paternal haplotype of Yao (middle), and CHM13 (bottom). Mitochondria DNA is not shown due to its short length relative to the large scale. Ancestry prediction (EAS, East Asian; EUR, European; SAS, South Asian; AMR, ad-mixed American; AFR, African) was overlaid with the density of SNPs against CHM13. All feature regions were lifted over to CHM13. The black vertical bars indicate the density of SNPs. **B.** Map of Hongtong migration routes in Ming Dynasty. Insert shows the sampling site of T2T-YAO and the adjacent TaoSi site—the ruin of Emperor YAO’ capital. **a.** sample site; **b.** TaoSi site. Map Content Approval Number, GS(2019)1371.

### Sample and sequencing

The goal of this work is to construct a complete and accurate reference genome for the Chinese Han population. Unlike hydatidiform mole cell lines, which have unstable genomes and a high chance of accumulating mutations and translocations during construction and passages as revealed by sequencing data in CHM13 (19), our study specifically collected a PBMC (Peripheral blood mononuclear cell) sample from a real person. This approach ensures greater accuracy and reliability in the generated reference genome. Furthermore, to achieve a more representative reference genome, this study specifically collected samples from Hongtong County, which is the starting point of the latest country-wide mass migration in China (**Figure 1B**). This migration, which destinations had involved 28.5% of counties in Ming territory, has had a significant impact on population distribution of their descendants throughout China and beyond, making it an ideal location for sampling. Additionally, the proximity of Hongtong to the ruins of the capital city where the legendary Emperor YAO lived over 4200 years ago (**Figure 1B**) also adds to its importance as a location for sampling. After this, we named the new reference genome “T2T-YAO”.

To construct the T2T diploid genome, we conducted a trio-based assembly by collecting multiple PBMC samples from the trio (son and both parents). Before sequencing, we conducted karyotyping test with 400 band on all three samples and ruled out any chromosomal disease in the family (**Figure S1**). This was followed by sequencing using multiple technologies, including: 92⨉HiFi sequencing from PacBio(20, 21), 336⨉ONT (Oxford Nanopore ultralong-read) sequencing, including 42⨉ super-long reads above 250kb(22), 584⨉Illumina Arima Genomics Hi-C sequencing(23), BioNano optical maps(24), and Illumina Hiseq 150bp for the son and parents (with 278⨉ and 116⨉ coverage, respectively). All the above sequencing coverage was calibrated in terms of an average haploid human genome, which is 3Gb. In pursuing high-fidelity and accuracy, we specifically selected long DNA fragments greater than 100kb for the library construction and applied the SUP model for the base-calling in the ONT sequencing(25). The final ONT sequencing data achieved an N50 of 128kb with a median base-calling accuracy of 98% for single reads.

### Diploid assembly, polishing, and assessment

To construct the T2T-YAO diploid genome, we followed a similar strategy to the assembly of CHM13 but phased the filial ONT reads according to the haplotype-specific markers identified by the Hiseq sequencing data of each parent. Then we constructed a haploid-resolved de Bruijn graph using HiFi reads (26) and simplified the assembly graph by iteratively integrating the phased ONT reads into it (27). The automated pipeline-verkko (https://github.com/marbl/verkko) has integrated these steps and assembled the sequencing reads of YAO into a diploid genome (maternal, 22+X+M; paternal, 22+Y), leaving only 90 gaps within the 46 chromosomes. Most of the gaps are due to large scale repeats, particularly in the centromeres, the Yq12 region, and the short arms of the acrocentric chromosomes 13, 14, 15, 21 and 22 (SAACs). Next, using only super-long ONT reads (>250kb) and HiFi reads with unique or low-frequency k-mers, we successfully closed all the gaps by extending the overlapping HiFi reads guided by phased ONT reads from the boundary of the gaps. Finally, we validated the telomere sequences at the ends of all the 46 chromosomes and confirmed no intervening model sequences or gaps, achieving the T2T assembly of YAO which includes the complete sequences of 44 autosomes, chromosome X, Y, and mitochondrion).

To improve the accuracy of the T2T-YAO genome, we conducted several major steps (**Figure S2**) modified from the polishing process for CHM13(28). First, Winnowmap2 (29) was used to map HiFi and ONT reads to the T2T-YAO genome in a repeat-sensitive manner. Next, pipelines of NextPolish2 (30), DeepVariant (31), and PEPPER-DeepVariant(32) were used to call SNV (single nucleotide variant)-like errors in a haplotype-sensitive manner, and Sniffles2 (33) to call SV (structural variations)-like errors. Finally, SNV-and SV-like errors were respectively filtered by Merfin (34) and Jasmine (35), and manually checked. This process identified a total of 32,861 and 29,087 SNV-like errors, eight and seven SV-like errors in the maternal (Mat) and paternal (Pat) haploid, respectively, which were corrected by BCFtools (36, 37). BioNano and Hi-C data were also used to identify potential SV-like errors, and only 28 discordant variations supported by ONT and HiFi reads were corrected manually. Ultimately, we obtained a fully-resolved diploid genome T2T-YAO with its total length of Mat and Pat haploid of 3,008,223,111 bp (3.01Gb) and 2,912,912,805 bp (2.91Gb), respectively. The size of both T2T-YAO haploids is comparable to that of T2T-CHM13, and the difference between the two haploids is mainly due to the different length of sexual chromosomes.

The highly polished T2T-YAO was then evaluated by Merqury (38), a reference-free assessment pipeline, using both Hiseq and HiFi reads to determine its completeness, assembly errors (**Figure 2A)**, and switching errors between haplotypes (**Figure 2B**). T2T-YAO reaches the reference-quality of accuracy with its quality value (QV) which represents a log-scaled probability of error for the consensus base calls reaching Q71.03 for Mat and Q71.52 for Pat haplotype **(Table 1)**. The quality is slightly less than the Q72.62 of the haploid genome-CHM13(28), but much higher than that of HG002 and CN1 (Q58.6∼60.1), the up-to-date highest-quality diploid human genome (11, 18). T2T-YAO achieved high quality, even after filling all the gaps, which process often introduces a large number of assembly errors. The QV scores of more than half of the chromosomes in T2T-YAO are greater than 75, and the QVs of chr5 in Mat and chr8 in Pat even reach infinity, which means no assembly error **(Table S1)**. The T2T-YAO shows comparable percentage of false duplications but much less collapsed repeats than HG002 **(Table 1)**. The copy numbers of several important multicopy genes in T2T-YAO, including the rDNA cluster, and the lengths of satellite regions in the X chromosome were estimated by performing droplet digital PCR (ddPCR) in comparison to *in silico* PCR (**Table S2**). All the assessments of T2T-YAO ensure that it is highly qualified as a reference genome.

**Figure 2.**
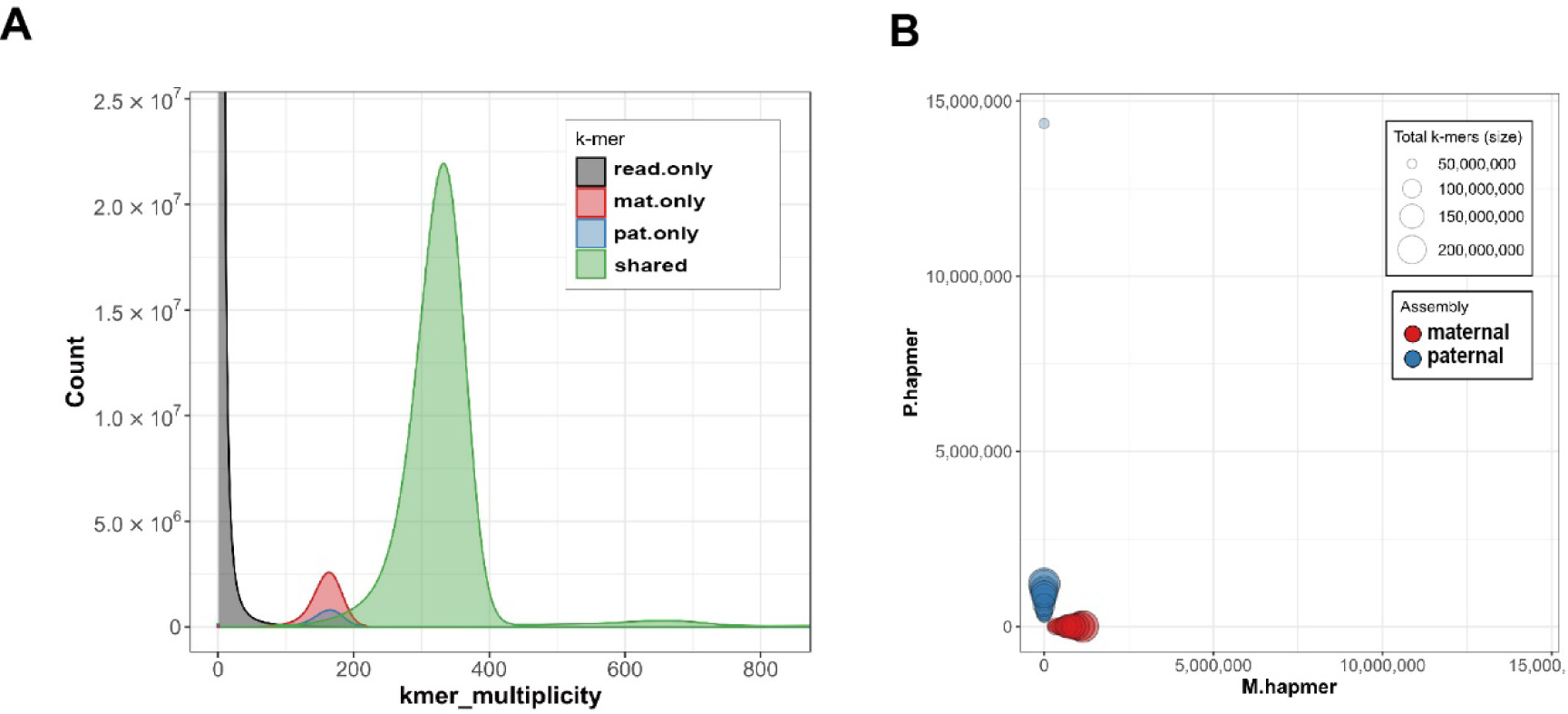
The completeness and quality of T2T-YAO assembly. **A**. Merqury assembly spectrum plots for the k-mer-based evaluation of assembly error and completeness. The distribution of k-mers present in the reads and maternal/paternal assemblies are separately colored. **B**. Hap-mer blob plot of the T2T-YAO assembly. Red blobs represent maternal chromosome; blue blobs are paternal chromosome. Blob size is proportional to k-mer size in each chromosome.

### The ancestry markers

The local ancestry inference based on random forest models (39) trained by the SNPs called from the 1000 Genome Project (KGP) dataset (40) demonstrates that the majority of genome YAO is of East Asia origin and admixed with sporadic predicted markers of South Asia, Europe, and America (**Figure 1A**). The positions of the ancestry admixture are not consistent between the two haplotypes due to the interindividual diversity in SNP profiles in Han population. The markers of South Asia are a little more than those of Europe and America in both haplotypes, suggesting more genetic exchange between the two Asia ethnic groups than with groups in other continents, which is consistent with previously reported study (41). When inferred with the same random forest model, the ancestry of CHM13 is mainly of European origin admixed with a few predicted markers of East Asia, South Asia, and Africa.

We further identified the Y-haplogroup of YAO using yHaplo (42), which is a tool to build a tree of phylogenetically significant SNPs accumulated in the non-recombining portion of the male-specific region (MSY)(43), and compared it to the Y-DNA Haplogroup tree in ISOGG (International Society of Genetic Genealogy) database(44). The Y-chromosome haplogroup of YAO is identified as O-F2137 (O2a2b1a1a1a2a), which is one of the main descendant groups of O-M122, the predominant Y-haplogroup in China (45, 46), and consistent with the established ancestry of YAO. Interestingly, O-F2137 haplogroup is also identified in one of the ancient DNA sample in Shimao Shengedaliang, a Neolithic site in Shenmu County, Shaanxi (47). The site is dated to ∼2000 BC and is located close to the Taosi site in Shanxi, the ruin capital of Emperor YAO, near the village where we sampled. This finding suggests a potential genetic continuity in the region dating back to the earliest days of human habitation in this part of China.

### Exclusive sequences and genes

We performed pairwise alignment among the three haplotypes (Mat and Pat in YAO and CHM13) using MUMmer (48) to get a full view of the sequence similarity among them. The dot-plot of the alignments exhibit a two-phase pattern for all pairings in term of the length and identity of alignment: one is of perfect alignments that longer than 50kb with high identity (mostly >99.5%); the other is of poor alignment less than 50kb with low identity (90-99.5%) (**Figure S3**). The perfect alignments indicate the orthologous regions that constitute the majority of the chromosomes, and the rest 280-350Mb sequences (∼10%) in each haplotype are not or poorly aligned to others, which mainly locate in the regions of heterochromosome, such as centromeres and the Yq12 region (**Figure 3A**). Heterochromosome regions, comprised of highly repetitive sequences, are more instable during chromosomal duplication and diverse among individuals than the other parts of the genome, which has been found to contribute to aging, neurodegeneration, and other disorders (8, 49, 50).

**Figure 3.**
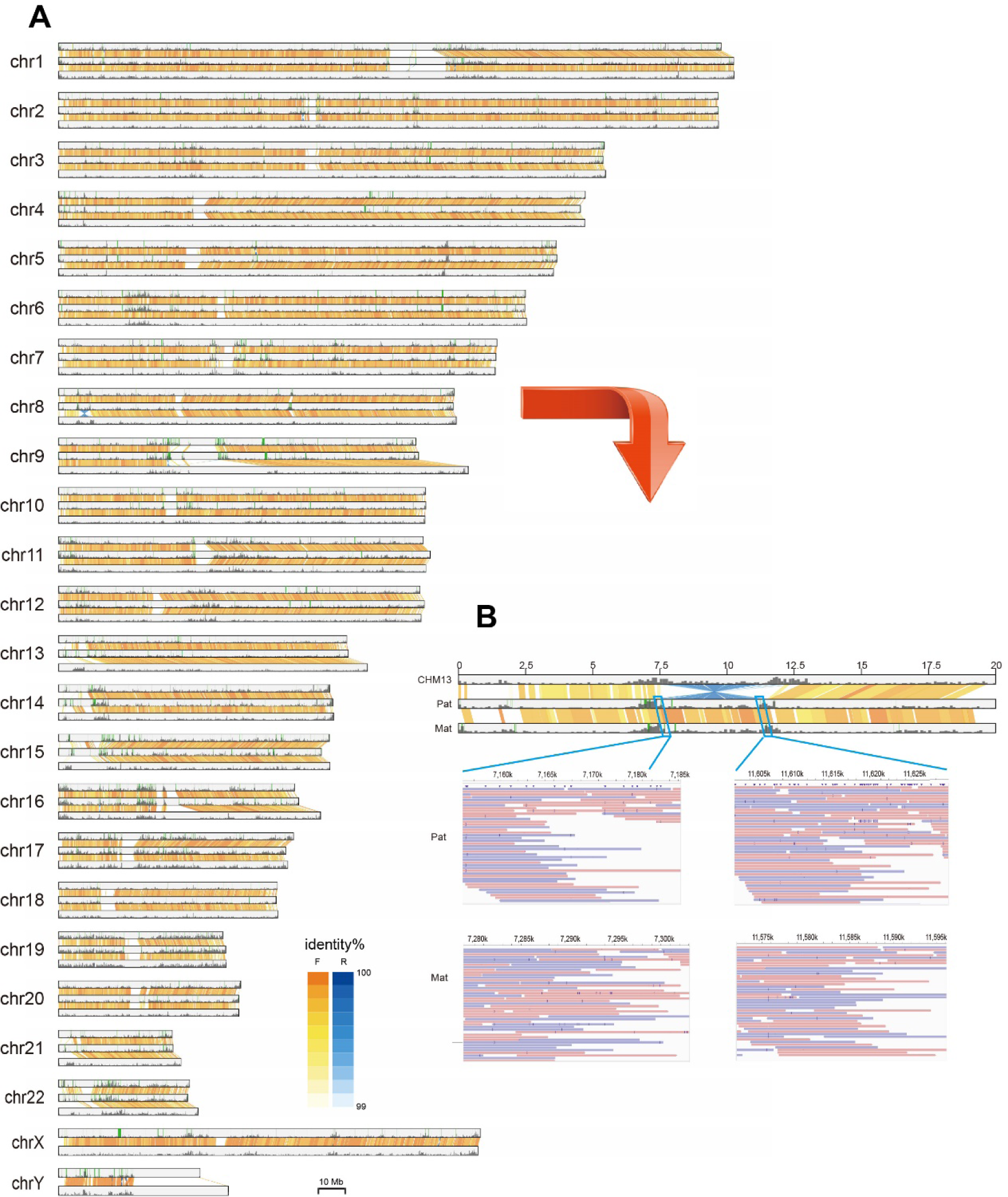
The overview of genomic variation between YAO and CHM13. **A**. The alignments between the chromosomes of maternal, paternal, and CHM13 (v2.0) haplotype. Each set of chromosome includes maternal haplotype of Yao (top), paternal haplotype of Yao (middle), and CHM13 (bottom). Alignments with length>100kb and identity > 98% are displayed by lines between the chromosomes. The color indicates the direction of alignment, orange, forward; blue, reverse; darkness indicates the identity. The black vertical bars indicate the density of genes with exclusive genes labeled by green lines. **B.** The upper panel is the alignment map of the distal region (0-25Mbp) of Chr8 short arm where there is a large inversion in both maternal and paternal haploid. Corresponding region in CHM13 is aligned on the top of the map for comprison. The bottom panel shows the Hifi reads mapped to the break points in the maternal and paternal assemblies.

It is notable that the total length in Mat and Pat haploids of poorly aligned sequences to CHM13 in the 22 euchromatic autosomes is 321.9 Mb and 329.2Mb, respectively, but reduce to 292.1 Mb and 299.8 Mb of them when aligned to each other, indicating better alignment between the Mat and Pat than that to CHM13 (**Table S3**). Furthermore, in the perfect alignments longer than 50kb, the weighted average identity between the two haplotypes of YAO is 99.94%, higher than that of 99.83% between YAO and CHM13 (**Figure 3A**). We then investigate the sequence similarity in the centromere regions among the haplotypes. The result shows some more alignments and higher similarity in the Mat vs. Pat than in YAO vs. CHM13 pairings (**Figure S4**). All these alignment results indicate that ∼10% of sequences in each haplotype are of unique and represents most of the inter-individual genome diversity. The haploids of YAO share more sequences and possess higher identity than comparisons to CHM13, implying a greater genomic distance between the ethnic groups compared to that within the Han population. Then we annotated the T2T-YAO genome and compared its gene components to CHM13, for which the Comparative Annotation Toolkit (CAT) (51) and Liftoff (52) were applied to project the GENCODE v43 annotation of reference GRCh38 (53) onto the T2T-YAO assembly. The annotation of the T2T-YAO genome was further refined by merging the annotations obtained from both tools, identifying 64,132 and 62,285 genes in Mat and Pat haploid, respectively, similar to the number of genes found in CHM13 (**Table 2**). Either Mat or Pat haplotype has ∼3000 exclusive genes other than CHM13, 2646 of which are shared in-between Mat and Pat haploids (**Figure S5)**. Furthermore, Mat and Pat haploids share a similar pattern of their exclusive genes dispersed along chromosomes, with some hotspots in the peri-centromere regions of low identity to CHM13 (**Figure 3A**), suggesting that haploids of Han origin are much closer to each other than to CHM13 in term of the genes they harbor.

Among the 2646 exclusive genes that is shared by both haploids of YAO, 232 are predicted to be protein-coding (**Table S4**), while the rest are noncoding. Mat and Pat haploid also has a few hundred exclusive genes to each other, including dozens of protein-coding genes (**Table S5-6**). According to their annotations, most of the exclusive protein-coding genes are pseudogenes, gained gene copies, or members of large gene family, such as the 30 genes in the Olfactory Receptor Family. Notably, 78 exclusive protein-coding genes are annotated as “novel protein” with unknown functions, and 76 out of them are single-copy ones. These novel proteins exclusive to CHM13 potentially highlight the unique features in the genomes of Han population in term of function and deserve further studies in the future.

### Genome variations in homologous regions

We first compared the small variations, including SNVs (single nucleotide variations) and small indels (<50bp), in the orthologous sequences among the three haploids, for which we used GRCh38 as the reference in order to accurately annotate the variations. A total of 3.03 Mb, 2.94Mb, and 2.92Mb of small variations are identified in Mat, Pat, and CHM13, respectively, largely comparable to those averagely identified in each haplotype in the pangenome studies (13, 14). When only taking into account the autosomes, the total length reduces to 2.92 Mb, 2.94 Mb, and 2.82 Mb, respectively, among which Mat and Pat share another 0.60 Mb small variations in-between besides the 0.88Mb shared by the three, but only share 0.366 Mb and 0.367 Mb with CHM13, respectively, suggesting a greater genetic distance between Han and European populations (**Figure S6**).

Most of the small variations locate in the intergenic region and introns, and only ∼0.05% small variations, mostly SNV, are in the CDS region of protein coding genes. Among the SNVs in CDS regions, more than a hundred are nonsense or frameshift mutations, which numbers are similar among the three haplotype and within the range of such mutations identified in the recent pangenome study(13) (**Table S7**). Interestingly, only 11 nonsense and 13 frameshift mutations are shared by both haplotypes of YAO, and all the affected genes are pseudogenes or members of gene family, implying that the loss-of-function mutations in them may not lead great damages to the host.

To investigate the overall genetic distance between the three haplotypes, we further compared their k-mer composite. As the k-mer composition varies between repetitive and non-repeat sequences, we first separated the repeat and non-repeat regions for each haploid genome. Then we calculated weighted dissimilarity matrix for non-repeat section and unweighted matrix for repeat section. In the PCoA plot, most homologous chromosomes are very close, indicating similar k-mer composition in each chromosome, except for chr1, 2, and the acrocentric chromosomes. These chromosomes showed a greater distance between CHM13 and the YAO haplotypes in both repeat and non-repeat sections (**Figure S7**).

Next, with T2T assembly of YAO, we are able to perform a more thorough analysis of the structure variations (SVs) among the three haplotypes in a pairwise manner. Using SVIM-asm (54), we inferred 26,701 and 28,642 SVs (>50bp) in Mat and Pat haploid, respectively, when compared to CHM13. Most of the SVs are indels in the range of 50~300bp, while the longer ones are sporadically distributed (**Figure S8**). SVs between Mat and Pat are fewer than their comparison to CHM13 in both term of number and total length by ∼10% (**Table S8**), indicating close genetic distance among the YAO’s haplotypes. We further investigated the larger SVs (> 100 kb) and identified 86 and 88 in Mat and Pat haploid, respectively, when compared to CHM13 using SyRI (55). A fraction of 82 and 81 SVs locate in the autosomes, a little more than the 76 SVs between Mat vs. Pat comparison. However, the inversions and translocations in YAO vs. CHM13 pairings are often much larger than those between Mat vs. Pat, making their total lengths several folds longer (**Table S9**).

The largest inversion we identified, which spans 4Mb in length, is located on the short arm of chromosome 8 (8p23) of both YAO haploids when compared to CHM13. Careful verification of the assembly in both Mat and Pat haploids shows strong support from multiple coverage of HiFi and ONT reads in comparison to the break points in CHM13 where are entirely devoid of reads (**Figure 3B, Figure S9**). Interestingly, the 8p23 inversion has also been reported as a structural polymorphism in previous genetic research (56) and in the recent-released high-quality genomes of Han Chinese individuals (17, 18). Near both sites of the break, we identified many exclusive member genes of the Olfactory Receptor Family unique to CHM13, however, the inversion does not disrupt or fuse any genes, although it does switch the intervening genes to the opposite strand of the chromosome.

### Homology mosaics in the short arms of acrocentric chromosomes

The assembly of the short arms of acrocentric chromosomes (SAACs) is a challenging task due to their highly repetitive sequences and extensive recombination among homologous and heterologous acrocentric chromosomes, such as the Robertsonian translocation between chr.14 and chr.21 observed in cytogenetic studies(57). The highly entangled contig components of SAACs in the assembly graph (**Figure 4A**) demonstrates the complexity of unraveling their complete sequences, which has remained unachieved by previously published human diploid genome, including those reported in the recent pangenome studies(13, 14). Relying on the deep coverage of our highly accurate ultra-long ONT reads and the clear phasing between Mat and Pat haplotypes, we have been able to successfully complete all ten SAAC regions in T2T-YAO. 4›Pairwise alignment of the complete SAACs revealed the presence of almost identical sequences across heterologous chromosomes, forming homology mosaics with large amount of inversions, duplications, and translocations, especially among chr.13, 14, 21, and 22 (**Figure 4B**), making them hotspots of chromosomal abnormalities usually seen in genetic disorders and tumors. We then investigate the k-mer composition of the ten SAAC regions, and their pairwise distance in PCoA plot indicates the homogeny among them, which cluster together. In contrast, the long arms of homologous chromosomes show almost identical position but far from the heterogenous one (**Figure 4C**). The completion of the diploid assembly of T2T-YAO has provided direct sequence-based evidence for the homology mosaics nature of SAACs, further confirming the active recombination events in these regions (58).

**Figure 4.**
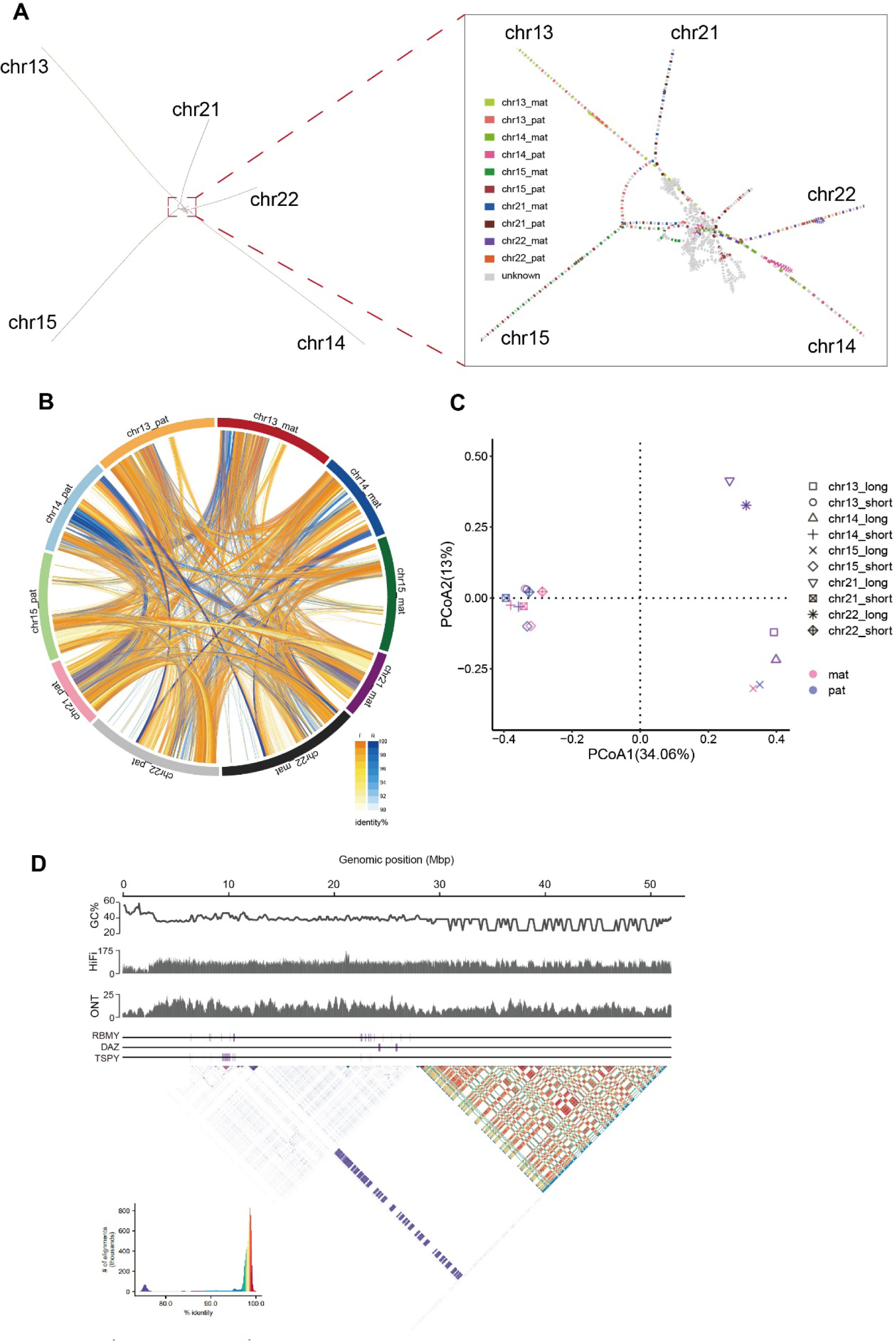
Assembly of the short arm of acrocentric chromosomes and chromosome Y. **A**. Bandage visualization of acrocentric chromosomes. Each pair of homologous chromosomes overlapped with each other and the short arms of the ten acrocentric chromosomes (SAACs) entangled together. Right panel shows the enlargement of the entangled short arms. The constitute block of chromosomes are colored according to the haplotype and chromosome it is assigned. **B.** Sequence similarity among the SAACs. The ten SAACs are permutated along the circumference. Pairwise Alignments between heterogenous chromosomes with length>10kb and identity> 90% are displayed by curves. The color indicates the direction of alignment, orange, forward; blue, reverse; darkness indicates the identity. **C.** The PCoA plot of the Jaccard distance among the short arms and long arms of the acrocentric chromosomes based on their k-mer composition. **D.** The structure of the complete chromosome Y-YAO. Peak charts show the GC content, coverage of the mapped HiFi reads and ONT reads (>100kb) along Y-YAO. The middle panel shows the positions of the ampliconic genes RBMY, DAZ, and TSPY on the chromosome. The bottom panel is the alignment dot plot of the tandem repeats in Y-YAO visualized by StainedGlass with sequence identity indicated by the color.

Each SAAC in T2T-YAO contains a locus of clustered rDNA genes, arranging in large stretches of tandem repeats without any rDNA cluster outside these regions, similar to CHM13. These rDNA clusters are highly homogenized, except for a few variations in the intergenic non-coding regions, rendering them susceptible to copy loss during chromosomal duplication(59). Being of paramount importance for cellular biology, the copy number of rDNA genes is tightly regulated to maintain homeostasis and is associated with the state of cell proliferation and DNA methylation (60, 61). The copy number of rDNA clusters in each chromosome is highly variable in YAO, and the total copy number in Mat and Pat haploids is 79 and 154, respectively, significantly fewer than the 209 copies in CHM13, but consistent with our ddPCR results (**Table S2**) and previous reports (59). The amplification of the rDNA gene in CHM13 is possibly an adaption to the rapid proliferation of the hydatidiform mole cells.

### The architecture of chromosome Y

The assembly of chromosome Y appears to be particularly challenging due to the high degree of repeat-sequence content, especially in the Yq12 region. The T2T assembly of our YAO-Y has a length of 51Mb, which is smaller than the T2T-Y in CHM13 v2.0 by about 10Mb (62), but still within the range of length polymorphism of chromosome Y reported recently, ranging from 45.2 to 84.9 Mb (63). The difference is primarily due to a contraction in the Yq12 heterochromatic region, which spans 24 Mb in YAO and is comprised of alternating human satellite 1 and 3 blocks, which are repetitive sequences with high identity. The correctness of our YAO-Y assembly is supported by the evenly covered HiFi and ONT reads mapping to it (**Figure 4D**), and some segments in the Yq12 region showing decreased HiFi coverage are composed of DYZ2 repetitive satellite sequences of high A/T content, which are known to be biased in HiFi sequencing(28) **(Figure 4D)**. The architecture of YAO-Y is consistent with previous report, containing the pseudo-autosomal regions (PARs) at both ends, X-transposed regions, ampliconic sequences, heterochromatic satellites, and X-degenerate regions. Most of the orthologous blocks aligned perfectly with those in CHM13-Y, with the exception of some translocations and inversions in the ampliconic region and the low identity region of Yq12 (**Figure 3A**). In YAO-Y, the copy numbers of the ampliconic genes of TSPY, DAZ, and RBMY are 53, 4, and 32 copies, respectively, consistent with those in CHM13-Y and confirmed by our *in silico* PCR and ddPCR results (**Table S2**). The ampliconic structures of these genes are also fully preserved in the ampliconic regions (**Figure 4D**), similar to the pattern observed in CHM13-Y(62).

### A truly complete diploid reference genome for Han population

T2T-YAO includes *de novo* telomere-to-telomere assembly for the diploid human genome of a single Han individual, with 44+XY chromosomes that contain 22 autosomes and X/Y in each Mat/Pat haplotype, inherited separately from the mother and father. This diploid assembly poses significant challenges due to the regions of high repetitiveness and extensive recombination, particularly in the SAAC regions. Despite these challenges, T2T-YAO has been completed thanks to the advent of the high accuracy long-reads sequencing technology and pipelines for reads phasing and diploid assembly. T2T-YAO achieved a comparable quality to T2T-CHM13 and no gaps filled with artificial sequences or model nucleotide, representing a truly complete and accurate genome sequence of a real individual, including the Y chromosome. The temporal-spatial haploid-phased expression of allele genes in various tissues is a critical issue in the biomolecular atlas of human body, with implications for understanding all kinds of biological processes, including embryo and organ development, aging, as well as disease pathogenesis. The differential role of allele genes in such processes is largely unknown, emphasizing the importance of building a diploid T2T reference for human genome. T2T-YAO and its iPS cell lines thus provide an essential resource for studying the cellular biological processes in a phased manner, including RNA transcribing and epigenetic modifications in DNA and RNA. Further characterization of the genetic and biological features of the T2T-YAO genome is ongoing, including complementing the annotations of its exclusive genes by transcriptome sequencing of the iPS cells and investigating the genomic diversity of Han population by variant calling against YAO.

The differences of both YAO haplotypes to CHM13 are much greater than their in-between differences in terms of ancestry markers, exclusive sequences and genes, k-mer composition, sequence-level and structure variations. The inter-ethnic difference, which have long been underestimated due to the preoccupation with short-read sequencing technologies, may contain significant genetic features associated with ethnic populations, diseases, and various phenotypes. The previously unknown distinct sequence and structure variations in YAO would undoubtedly provide valuable insights and further our understanding of human genomics and its applications in clinical medicine. The availability of T2T-YAO would of course improve mapping and analysis of short-read and long-read data from samples of Han individuals, highlighting the necessity of building a special T2T reference for the huge population. Furthermore, the comprehensive annotation of T2T-YAO is ongoing, providing a valuable resource for researchers to study the genetics and biology of the genome of Han people, improving the practice in health research and precision medicine in China.

## Materials and Methods

### Sample collection and Ethics

Application of the study was submitted and approved by the Ethical Review Committee of Linfen Central Hospital (No. 2022-20-1). Collection and storage of human samples were registered at and approved by the Human Genetic Resources Administration of China (HGRAC). Written informed consents were obtained from the participants. For the YAO genome, the fresh blood sample was collected from a healthy male of northern Han and both of his parents. Routine blood test and liver and kidney function were conducted to exclude the potential diseases.

### Karyotyping and iPS cell line

For karyotype analysis, peripheral blood lymphocytes (PBLs) were cultured. The fresh and anticoagulated blood samples were inoculated in the culture bottles, and then the culture bottles were placed in an incubator set at a constant temperature of 37°C for 68 hours. Prior to harvesting the PBLs, colchicine was added into culture medium for an additional 60 minutes of cultivation. Then the PBLs were harvested, and treated with KCl solution followed by fixation with Carnoy’s fixative (Methanol: Acetic Acid = 3:1). Chromosomes were stained using Giemsa, and images of chromosome spread were captured using an automated cytogenetic imaging system (Leica, GSL-10).

Peripheral blood mononuclear cell (PBMCs) were obtained from peripheral blood with Ficoll-Hypaque density gradient centrifugation and expanded in DMEM supplemented with 10 % fetal bovine serum, 100 units/ml penicillin, and 100 μg/ml streptomycinin a humidified incubator with 5 % CO2 at 37 ◦C. The PBMCs were reprogrammed by using Sendai virus (CytoTuneTM-iPS 2.0 Sendai Reprogramming kit, Thermo Fisher Scientific) according to the manufacturer’s protocol, which includes four Yamanaka factors, OCT4, SOX2, KLF4 and C-MYC, that are sufficient for efficient reprogramming. Subsequently, the transfected iPSCs were suspended in Essential 8™ (Thermo Fisher Scientific, A1517001) medium on 0.1 mg/mL Matrigel (Corning, 354277)-coated plates and incubated at 37 ◦C in 5 % CO2 atmosphere. From the next day, the erythroid medium was half-changed with ReproTeSR (StemCell Technologies) medium every other day. 14 days after transfection, iPSC colonies were picked up and cultured in MTeSR PLUS medium (StemCell Technologies, 05825) in vitronectin (1:100 dilution, GIBCO, A14700) coated 6-well plates. Cells were passaged using Versene (GIBCO, 15040-066) and MTeSR PLUS according to the manufacturer’s instruction at a ratio of 1:10.

### Genomic DNA extraction

High molecular weight genomic DNA was prepared by the CTAB method and followed by purification with QIAGEN® Genomic kit (QIAGEN, 13343,). Ultra-long DNA was extracted by the SDS method without purification step to sustain the length of DNA. DNA purity was detected using NanoDrop™ One UV-Vis spectrophotometer (Thermo Fisher Scientific, USA), DNA degradation, and contamination of the extracted DNA was monitored on 1% agarose gels. At last, DNA concentration was further measured by Qubit® 4.0 Fluorometer (Invitrogen, USA).

### Ultra-long Nanopore Sequencing

For each ultra-long Nanopore library, approximately 8-10 μg of gDNA was size-selected (>50 kb) with SageHLS HMW library system (Sage Science, USA), and processed using the Ligation sequencing 1D kit (SQK-LSK109, Oxford Nanopore Technologies, UK) according the manufacturer’s instructions. About 800ng DNA libraries were constructed and sequenced on the Promethion (Oxford Nanopore Technologies, UK) at the Genome Center of Grandomics (Wuhan, China).

### PacBio HiFi sequencing

For PacBio HiFi whole-genome sequencing, SMRTbell target size libraries were prepared according to PacBio’s standard protocol (Pacific Biosciences, CA, USA) using 15kb preparation solutions. The main steps include (a) gDNA shearing by g-TUBEs (Covaris, USA), (b) DNA damage repair, end repair and A-tailing, (c) ligation with hairpin adapters from the SMRTbell Express Template Prep Kit 2.0 (Pacific Biosciences), (d) nuclease treatment of SMRTbell library with SMRTbell Enzyme Cleanup Kit, (e) size selection by the BluePippin (Sage Science, USA), and binding to polymerase. The SMRTbell library was then purified by AMPure PB beads, and Agilent 2100 Bioanalyzer (Agilent technologies, USA) was used to detect the size of library fragments. Sequencing was performed on a PacBio Sequel II instrument with Sequencing Primer V2 and Sequel II Binding Kit 2.0 in Grandomics.

### Illumina short-read sequencing

For Illumina sequencing, the DNA of offspring and his parents were sequenced for NGS. 1μg genomic DNA was randomly fragmented by Covaris, and then the fragments were selected by Magnetic beads to an average size of 200-400bp. Eligible fragments were end repaired and then 3’ adenylated. Adaptors were ligated to the ends of these 3’ adenylated fragments, which used to amplify the fragments. The PCR products were heat denatured, and circularized by the splint oligo sequence. Ultimately, the single strand circle DNA (ssCir DNA) were formatted as the final library. The library was amplified with phi29 to make DNA nanoball (DNB), then DNB were load into the patterned nanoarray, and pair end 100/150 bases reads were generated in the way of combinatorial Probe-Anchor Synthesis (cPAS).

### Hi-C sequencing

For Hi-C sequencing, purified DNA were digested with 100 units of DpnII and incubated with biotin-14-dATP. The ligated DNA was sheared into 300−600 bp fragments, and then was blunt-end repaired and Atailed, followed by purification through biotin-streptavidin-mediated pull down. Finally, the Hi-C libraries were quantified and sequenced using the Illumina Novaseq/MGI-2000 platform.

### Bionano

For Bionano analysis, extracted DNA were subject to manufacturer recommended protocols for library preparation (Bionano PrepTM Animal Tissue DNA Isolation Kit (CAT#80002) and optical scanning provided by BioNano Genomics (https://bionanogenomics.com), with the labeling enzyme Direct Label Enzyme(DLE) (Bionano PrepDLS Labeling DNA Kit, CAT#80005). Labelled DNA samples were loaded and run on the Saphyr system (BioNano Genomics) in Grandomics.

### Genome assembly

Before assembling, Hiseq reads of the offspring and both parents were trimmed and only bases with Phred Quality Score >30 were retained for the following k-mer calling. Hapmers from each parent were input to verkko (v1.0) along with the HiFi and ONT reads, and the initial assemblies were constructed with verkko in the trio mode. The output fully-phased contigs of Mat and Pat genome of YAO were further scaffolded guided by CHM13. To fill the gaps, ultra-long ONT reads >250kb and HiFi reads with low-frequency k-mers were mapped to the extracted 5 kb of flanking regions, and close the gap iteratively extension the aligned reads. These alignments were further visualized and manually examined. All the final chromosome were checked for telomere sequences at both ends to ensure its completeness and free of any model nucleotides.

### Polishing and validation

To correct potential single nucleotide variants (SNV) and structural variants (SV), we followed the polishing pipeline of Mc Cartney, et al (28). Long reads from the offspring are binned into paternal and maternal groups based on the presence of the haplotype-specific k-mers by TrioCanu(64). The alignment was conducted between corresponding haplotype and reads using Winnowmap2(29). DeepVariant(31) and PEER-DeepVariant(32) were used to call small errors from self-alignments of HiFi and ONT reads, respectively. Similarly, calls with low allele fraction support or low genotype quality (GQ) score were removed (GQ < 30 for the HiFi calls and GQ < 25 for ONT SNP calls, and variant allele frequency (VAF < 0.5)). The remaining SNVs were merged using the custom script(28) and then Merfin(34) were used to ensure that no new false kmers were introduced due to the correction of the SNVs. The SVs were detected by Sniffles(33) from the alignments of ONT and HiFi, and finally SVs with supportive reads less than 60% were removed. Jasmine(35) was used to merge the SVs. The SVs identified by both of ONT and HiFi reads were manually inspected and validated in Integrative Genomics Viewer (v2.6) and the true SVs were corrected.

### Assessment

Completeness analyses for the diploid assemblies was calculated with BUSCO v3.1.0(65) using the mammalia_odb10 lineage set (https://busco-archive.ezlab.org/v5/). 21-kmers of reads were collected from Illumina and HiFi reads of offspring and the parental Illumina reads using Meryl, and Merqury(38) was used to calculate QV, completeness and phasing statistics. Collapsed and expandable sequences were calculated using SDA(66). Segmental duplications (SDs) were detected by BISER(67).

### Droplet digital PCR

Genomic DNA was isolated from the iPS cell line using the Magetic Univeral Genomic DNA kit (TIANGEN, DP705-01). Copy numbers of ampliconic genes were validated through droplet digital PCR (ddPCR). Primers, gDNA concentrations, and restriction enzymes are referred to previous study(4, 68). ddPCR reactions were performed using the dye method ddPCR detection kit (TargetingOne) and TargetingOne ddPCR system. Briefly, each reaction consists of 15 uL 2x ddPCR Supermix, 0.2 uL of restriction enzyme for fragmentation, 3 uL 10 uM primer mix, 1 uL of 2-50ng DNA template and 10.8 uL with nuclease free water. Mastermixes were then emulsified with 180ul droplet generator oil using a droplet generator according to the manufacturer’s instructions. After droplet generation, thermocycling was performed with the following parameters: 5min at 37°C, 10 min at 95°C, 45 cycles consisting of a 30-sdenaturation at 94°C and a 60-s extension at 55°C, followed a hold at 12°C for 5 minutes. Control reactions without the DNA were performed to rule out non-specific amplification.

### Gene annotation

The final annotation combined the Comparative Annotation Toolkit (CAT) result and the Liftoff annotation. First of all, alignment to GRCh38 was generated with Cactus(69) and used as input for CAT(51) along with the GENCODEv43 annotation. Liftoff(52) was run to map genes from the Gencodev43 to the ChTY001. Predictions of Liftoff that did not overlap any CAT annotations were added using Bedtools intersect. Common repeat elements were identified by RepeatMasker (v4.1.0).

### Ancestry analysis

RFMix 2(39) (https://github.com/slowkoni/rfmix) was used to infer the local ancestry of the ChTY001 genome. 2531 individuals with 30× sequenced in the Phase 3 release of the 1000 Genomes Project were used as a set of reference samples for ancestry (https://www.internationalgenome.org/data-portal/sample)(70). Biallelic SNVs of against the GRCh38 reference were obtained from 1000 Genomes Consortium (http://ftp.1000genomes.ebi.ac.uk/vol1/ftp/data_collections/1000_genomes_project/re lease/20181203_biallelic_SNV/). Genetic map for GRCh38 was obtained from Beagle (http://bochet.gcc.biostat.washington.edu/beagle/genetic_maps/). ChTY001 (paternal and maternal assembly) and CHM13v2.0 variants were called on GRCh38 with dipcall (https://github.com/lh3/dipcall)(71). RFMix2 was performed with –c 50 –s 5, grouping the 1000 Genomes reference panel into superpopulations (African, Ad Mixed American, East Asian, European, Southeast Asian). For the chrX, we only used SNVs in pseudoautosomal regions (PARs) and female individuals in the reference panel for ancestry analysis. The computed ancestry regions were lifted over from GRCh38 to CHM13v2.0 using liftOver (https://genome.ucsc.edu/cgi-bin/hgLiftOver).

### Y haplogroup identification

ChTY001 chrY was aligned to hg19 ChrY sequence with dipcall (https://github.com/lh3/dipcall)(71) to identify SNPs. We used yhaplo v1.1.2 to build a tree and determine the haplogroup (https://github.com/23andMe/yhaplo).

### Genomic variation

To call the full spectrum of heterozygosity among CHM13, and the ChTY001 diploid genomes, we directly compared these three assemblies using Mummer (v4.0.0rc1)(72). Sequences of unalignment and SNP were generated by ‘delta-filter –m –i 99 –l 100000’ and followed by ‘dnadiff’. The SnpEff v5.0 program(73) was adopted to infer functional annotation of any SNPs or small indels (<50bp) and any potential deleterious effect on protein structure.

SyRi (v1.5) (74) was used to detect SVs longer than 100kb, including inversion, translocation and duplications from Mummer alignments. Svim-asm was used to detect the SVs longer than 50bp.

## Acknowledgments

This study was supported by the grants of Linfen soft science research project (2126), National and Provincial Key Clinical Specialty Capacity Building Project 2020 (Department of the Respiratory Medicine), Peking University People’s Hospital Scientific Research Development Funds (RDY2022-11), the National Science Foundation of China (31970568), and the National Key Research and Development Program of China (2021YFC2301000).

## Author contribution

YKH, YNC, SMG, JH, RL, and YLZ carried out the assembly, polish, gene annotation, and drafted the manuscript. YKH, XQM, ZLD, LLZ, WYY, JBX, WJB, FFY, XC, and PAZ participated in the sequence alignment and manual correction. RHW, YFM, CJS, JC, JW, JWL, JW, XYH, QYL, MZJ, HLY, SXS carried out the validation, segmental duplications and data generation. GYL, YW, YX, YLM, YWC, and KQW participated in the repeat annotation and the draft editing. JB, WX, FW, WTN, MQZ, YY, and SNL participated in the variants and the supplement. YK and ZCG conceived of the study, participated in its design, coordination, supervision, the assembly, and helped to draft the manuscript. All authors read and approved the final manuscript.

## Tables

Table 1. Comparison of the assembling quality and repeat annotations among T2T-Yao, T2T-CHM13(v1.1), and HG002

Table 2. Comparison of the gene annotations between T2T-YAO and CHM13v2.0

## Supplement Figures

Figure S1. Karyotype of samples.

Figure S2. Polishing steps of T2T-YAO assembly.

Figure S3. Dot-plot of the alignments between Yao and T2T-CHM13v2.0. Each point represents an alignment. A, compare Yao.mat to CHM13v2.0; B. compare Yao.pat to CHM13v2.0.; C. compare Yao.pat to Yao.mat; D. compare Yao.mat to Yao.pat.

Figure S4. The similarity between YAO and CHM13v2.0 centromere regions. Each set for chromosome (chr) from top to bottom was Yao.mat, Yao.pat, and CHM13V2.0. Regions with sequence identity over 98% and longer than 100kbps are connected and colored by alignment direction. Orange, the join is forward; blue, reverse.

Figure S5. The Venn plot of gene/transcripts sharing between YAO and CHM13v2.0. noXY, genes or transcripts in chrX/Y were excluded from panels.

Figure S6. The Venn plot of small variants between T2T-YAO and CHM13. A. small variants shared in total chromosomes; B. small variants shared in autosomes.

Figure S7. The PCoA plots were drawn with k-mers shared in YAO and CHM13v2.0 assembly with/without repeat regions according to their presence/absence based on the Jaccard distance or their abundance based on Bray Curtis distance.

Figure S8. The type and length distribution of SVs between T2T-YAO and CHM13, and between two haploids.

Figure S9. The alignment of Chr8 (site 0-25Mbp) of maternal vs paternal assembly and paternal vs CHM13v2.0 assembly. Chromosomes from top to bottom were Yao.mat, Yao.pat, and CHM13v2.0. Hifi reads alignment to Chr8 of CHM13v2.0 is below chromosomes. Orange, the join is forward; blue, the join is reverse.

## Supplement Tables

Table S1. QV of total chromosomes of YAO assembly.

Table S2. Copy number validation of important multicopy genes in T2T-YAO with ddPCR.

Table S3. unaligned bases in T2T vs YAO.

Table S4. Protein-coding genes shared by maternal and paternal assembly and exclusive to Yao

Table S5. Maternal exclusive protein-coding genes of T2T-YAO assembly

Table S6. Paternal exclusive protein-coding genes of T2T-YAO assembly

Table S7. Small variations (<50bp) in T2T-YAO and CHM13v2.0 assembly

Table S8. the larger SVs (> 100 kb) between T2T-YAO and CHM13v2.0 assembly

